# Single-cell analysis suggests coordinated immune-cell redistribution between blood and cerebrospinal fluid in early multiple sclerosis

**DOI:** 10.64898/2026.08.04.742373

**Authors:** Nils Hallén, Sunjay Jude Fernandes, Soudabeh Rad Pour, Alexandra Gyllenberg, Yanan Han, Nicolas Ruffin, Narsis Kiani, Maria Needhamsen, Fredrik Piehl, Ingrid Kockum, Faiez Al Nimer, David Gomez-Cabrero, Jesper Tegnér, Maja Jagodic

## Abstract

Despite immune-cell infiltration being a hallmark of multiple sclerosis (MS), the interplay between the periphery and central nervous system immune responses is still incompletely characterized. We performed single-cell transcriptomic and V(D)J sequencing of paired blood and cerebrospinal fluid (CSF) immune cells from treatment-naive relapsing-remitting women with MS and compared them to age– and sex-matched healthy controls. Across major immune lineages, we identified coordinated, compartment-specific immune alterations, with enrichment of activated and memory lymphocyte populations in the CSF and concomitant depletion of related populations in peripheral blood, suggesting their recruitment from blood to CSF. Clonally expanded CD4 memory T-cells, together with activated, expanded IgM-positive B-cells, accumulated predominantly in the CSF of people with MS compared to healthy subjects. Interestingly, tissue-primed cytotoxic populations and CXCR3-associated memory populations were depleted from the CSF of MS, implying their recruitment to the target tissue in early disease. These findings reveal coordinated, compartment-specific immune changes in early MS and provide a systems-level view of immune-cell trafficking between peripheral and central nervous system compartments.

## Introduction

Multiple sclerosis (MS) remains one of the most common causes of neurological disability in young adults^1^. The disease pathology is characterized by an influx of immune cells into the central nervous system (CNS) and progressive myelin degeneration, leading to axonal loss and functional impairment^1^. Most MS patients are diagnosed with relapsing-remitting MS (RRMS), characterized by episodes of destruction followed by full or incomplete repair, and the majority eventually develop secondary-progressive disease^1^. However, inflammatory and progressive features co-exist at diagnosis^2^, and both contribute to disease progression^3^.

People with MS (pwMS) exhibit regional inflammatory plaques characterized by lymphocyte infiltration and diffuse changes. The strong genetic association with the HLA-DR15 haplotype^4^, together with adoptive-transfer experiments demonstrating that myelin-reactive T-cells can induce demyelinating disease^5^, have established autoreactive CD4 T-cells as key-players in the immune dysregulation of MS. Recent success of anti-CD20 therapies^6–8^ has generated renewed interest in B-cells, demonstrated to have the ability to present antigens and activate autoreactive CD4 T-cells^9^ and contribute to the pro-inflammatory environment, with an increased production of TNF, IL-6 and GM-CSF in pwMS^10,11^. Moreover, B-cells constitute the primary reservoir of Epstein-Barr virus (EBV), recently implicated as a necessary factor for MS development^12^.

Single-cell transcriptomic profiling has advanced our understanding of immune dysregulation in MS, utilizing cerebrospinal fluid (CSF) as a proxy of the inflammatory environment of the CNS^13–20^. Numerous studies have characterized the cellular composition of CSF in pwMS, consistently reporting expansions of class-switched memory and plasmablast-like B-cells, as well as activated CD4 and CD8 T-cell populations^13–15^. These studies have further identified transcriptionally distinct myeloid and microglia-like populations in the CSF of pwMS, including subsets enriched for antigen presentation and type I interferon response programs^16^. However, the majority of these investigations have focused exclusively on the CSF compartment, limiting the ability to place these findings in the context of systemic immune dynamics or to determine whether such cell states originate in, or are shaped by, peripheral immune responses. In contrast, relatively few studies have performed paired profiling of peripheral blood and CSF within the same individuals. Existing paired datasets have largely emphasized differences in cellular composition, for example, enrichment of follicular helper T-cells (Tfh) and B lineage populations in the CSF of pwMS, or have described broad inflammatory axes such as cytotoxicity and interferon signaling across compartments^14,15^. Recent work by Jacobs *et al.* performed the largest profiling of intrathecal immunity to-date, describing global neuroinflammatory features agnostic to disease^17^. They describe shared clonal pools between blood and CSF, and describe CSF-specific genetic regulation of expression. Furthermore, individuals with non-inflammatory neurological diseases are largely used as the control cohort in the majority of published studies. As far as we are aware, to date there exists no analysis of the entire immunological population of paired peripheral blood and CSF compartments in pwMS compared to matched healthy controls. As a result, a detailed understanding of how specific immune cell states and clonal populations are coordinated between the periphery and the central nervous system remains incomplete.

Here, we report single-cell transcriptomic and V(D)J analysis of fresh paired peripheral blood and CSF samples from newly-diagnosed untreated pwMS and age-matched healthy females.

## Methods

### Donor recruitment and ethical approval

CSF and blood samples were collected from treatment-I patients newly diagnosed with MS according to the 2017 McDonald criteria, following a first demyelinating event with short disease history. All patients were female and between the ages of 25 and 40 (**Table 1**). Healthy donors were sex– and age-matched (within a 5-year interval) with pwMS. Ethical approval (2009/2107-31, last amendment 2025-04961-02), was obtained from the Swedish Ethical Review Authority in Stockholm, Sweden (Regionala etiksprovnämden I Stockholm).

**Table 1.** Cohort characteristics.

|  | **MS**<br>(N=5) | **HC**<br>(N=4) | **NINDC**<br>(N=1) | **p-value** |
| --- | --- | --- | --- | --- |
| Age, mean (range) | 28.8 (24-34) | 29.8 (23-37) | 31.0 (31) | 0.8 |
| Female, N (%) | 5 (100) | 4 (100) | 1 (100) |  |
| Optic neuritis, N (%) | 1 (20) | 0 (0) | 0 (0) |  |
| OCB, N (%) | 5 (100) | 0 (0) | 0 (0) | **0.008** |
| Relapse, N (%) | 3 (60) | n/a | n/a |  |
| EDSS, median (range) | 1.5 (1.0-3.5) | n/a | n/a |  |
| Lesions number (MRI) |  |  |  | **0.016** |
| 1 – 9, N (%)<sup>1</sup> | 1 (20) | 0 (0) | 0 (0) |  |
| 10 – 20, N (%) | 2 (40) | 0 (0) | 0 (0) |  |
| > 20, N (%) | 2 (40) | 0 (0) | 0 (0) |  |
| Albumin (S), g/L, mean (SD) | 47.5 (6.0) | 45.5 (3.7) | 45.0 (n/a) | 0.8 |
| Albumin (CSF), mg/L, mean (SD) | 160 (21) | 160 (11) | 242 (n/a) | 0.9 |
| Mononuclear cells (CSF), cell/uL, mean (SD) | 5.00 (4.24) | 0.00 (0.00) | n/a | 0.2 |
| CXCL13 (CSF), ng/L, mean (SD) | 46 (29) | 2 (2) | 16 (n/a) | 0.077 |
| NfL (CSF), ng/L, mean (SD) | 868 (485) | 360 (106) | 17.240 (n/a) | 0.057 |
| κ-FLC (P), mg/L, mean (SD) | 14.5 (7.4) | 10.2 (2.0) | 18.0 (n/a) | 0.5 |
| κ-FLC (CSF), mg/L, mean (SD) | 4.66 (4.10) | 0.07 (0.03) | 0.13 (n/a) | **0.05** |
| IgG-index, mean (SD) | 1.06 (0.63) | 0.50 (0.03) | 0.50 (n/a) | **0.019** |
| Treated individuals, number (%) | 0 (0) | 0 (0) | 0 (0) |  |

N, number; MS, multiple sclerosis; HC, healthy controls; NINDC, non-inflammatory neurological disease control (mild head trauma); OCB, oligoclonal bands; EDSS, expanded disability status scale; MRI, magnetic resonance imaging; S, serum; CSF, cerebrospinal fluid; P, plasma; SD, standard deviation; <sup>1</sup>the actual range of 3-5. Statistical comparisons between MS vs. HC, Wilcoxon-test for numeric data, Fisher's exact test for categorical.

### Single-cell sequencing

Single-cell suspensions of CSF and peripheral blood mononuclear cells (PBMC) were loaded onto the 10x controller (10x Genomics). Transcriptomic and VDJ-libraries were prepared according to the manufacturer’s instructions. Libraries were sequenced on Illumina and Miseq platforms. Additional details can be found in **Supplementary Methods**.

### Single-cell sequencing analysis

Sequencing data was aligned using the *CellRanger* pipeline to GRCh38. Downstream transformations of data and statistical tests were performed in Python and R. Visualizations were generated in R. Additional details can be found in **Supplementary Methods**.

### Visualization portal

We generated an online portal for interactive exploration of expression data: https://single-cell-portal.jagodiclab.com (*to be released upon internal testing*).

### Data availability

Data will be deposited to the European Genome-phenome Archive (EGA).

## Results

### Global coordinated immune dysregulation in blood and CSF of pwMS

We performed transcriptome and V(D)J sequencing of fresh paired CSF and PBMC samples from young untreated women with MS, recruited during the initial diagnostic work up, and age-matched female controls (n_MS_=5, n_HC_=5, **Table1**), and we created an online portal for a data query (**Figure 1A**). We generated a dataset comprising 68,320 cells after quality control (QC), 40,159 from PBMC (mean n_HC_=3,539, n_MS_=4,498 cells) and 28,161 from CSF (mean n_HC_=1,240, n_MS_=4,569). After sample QC and batch-effect removal, cells were annotated into major lineages and further sub-clustered into 36 cell types (**Supplementary Methods**, **Supplementary Figure 1**). Notably, we addressed several aspects that have not been previously studied in MS. In particular, we made the distinction between shared-effector memory subsets, found in both blood and CSF, and tissue-restricted subsets, exclusively found in the CSF (**Supplementary Figure 2-6**). We also implemented a more recent classification of NK cell subsets^21^.

**Figure 1.**
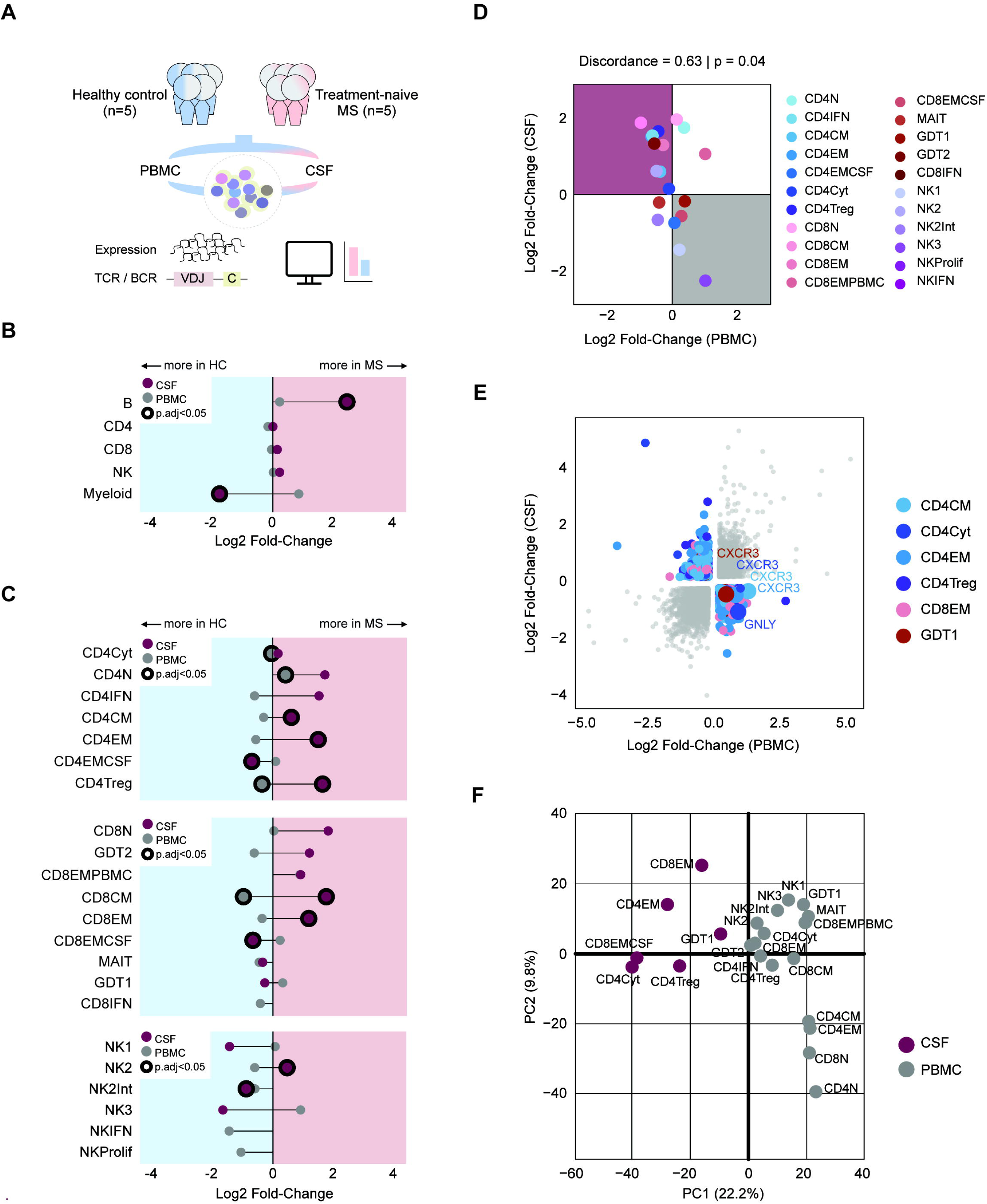
**A**. Schematic overview of experimental design. **B.** Relative log2 fold-change of lineage proportions in each tissue compartment across disease. Significant values are denoted with a black outline. **C.** Relative log2 fold-change of sub-cluster proportions inside each lineage across disease, segregated by tissue compartment. Significant values are denoted with a black outline. **D.** Log2 fold-change of sub-cluster proportions across disease in CSF (y-axis) or PBMC (x-axis) across disease. Highlighted quadrants illustrate discordant changes up in CSF (red) or up in PBMC (grey). **E.** Log2 fold-changes of differentially expressed genes across disease in each sub-cluster, CSF (y-axis) and PBMC (x-axis). Highlighted top genes by magnitude of difference between expression in each compartment. **E.** Principal component (PC) analysis of differentially expressed genes across disease for sub-clusters, segregated by tissue. Colored by tissue of origin.

On the lineage level, we observed a significant enrichment of B-cells and depletion of myeloid cells in the CSF of pwMS compared to matched healthy controls (**Figure 1B**). In contrast, no differences in abundances of CD4, CD8 or NK cells were found on the lineage level in the CSF or periphery. We next investigated potential perturbations within each of the three lineages, and we observed a significant increase in the relative proportion of central memory T-cells (CD4CM and CD8CM) in the CSF of pwMS (**Figure 1C**). Interestingly, in both CD4 and CD8 lineages, there was a significant increase in the effector memory T-cell subset shared between the CSF and blood (CD4EM and CD8EM) in the CSF of pwMS, accompanied by a significant decrease in the CSF tissue-restricted subset (CD4EMCSF and CD8EMCSF). A similar significant increase in the relative proportion of NK2 cells and a decrease of NK2-intermediate (NK2Int) cells was detected in the CSF of pwMS. Finally, our data corroborate, now in untreated pwMS compared to matched healthy controls, previously reported increase of CD4 regulatory T-cells (CD4Treg) in the CSF of pwMS^13–15^, and we further show a concomitant decrease in the periphery (**Figure 1C**).

We observed that many sub-clusters, albeit not always statistically significant, exerted a discordant relationship, i.e., relative proportions across disease differed in directionality between the periphery and CSF (**Figure 1C**). Thus, we leveraged the paired nature of the samples to demonstrate that nearly 70% of cell types exhibited such a discordant relationship across the compartments (p = 0.04, **Figure 1D**). We next addressed potential transcriptional discordance across the compartments by performing differential expression between pwMS and HC for each cluster. While we were limited due to constraints of differential testing in both compartments, we observed a number of genes that behaved in a discordant manner in several cell types (**Figure 1E**). Of interest, *CXCR3*, a brain-homing receptor, was found to be elevated in CD4CM, CD4EM and CD4Cyt in the periphery of pwMS, while it was decreased in CSF. This suggested a distinct MS response in blood and CSF, which we explored by performing Principal Component Analysis (PCA) on the results of differential expression. Strikingly, there was a strong axis of variation in the differentially expressed genes corresponding to the compartment, with genes like *CD74*, *SAT1* and *SNHG12* driving this separation (**Figure 1F**).

These results highlight global immune changes in early MS that affect both the CSF and the periphery, characterized by distinct transcriptional responses and a discordant relationship across the compartments indicative of active lymphocyte recruitment to the CNS.

### Tissue-restricted lymphocytes are depleted from CSF of pwMS

We aimed to better characterize the T-cell and NK lineage (**Supplementary Figure 2-4**), which exerted a discordant relationship across the compartments in pwMS (**Figure 1C, D**). We observed striking compositional differences between CSF and blood, regardless of disease, with effector memory cells, in particular tissue-restricted subsets (CD4EMCSF and CD8EMCSF), comprising a major portion in the CSF, while naive cells (CD4N and CD8N) were the most abundant in the periphery (**Figure 2A**). The CSF NK lineage largely consisted of classical and intermediate NK2 cells, while NK1 and NK3 dominated blood samples (**Figure 2A**).

**Figure 2.**
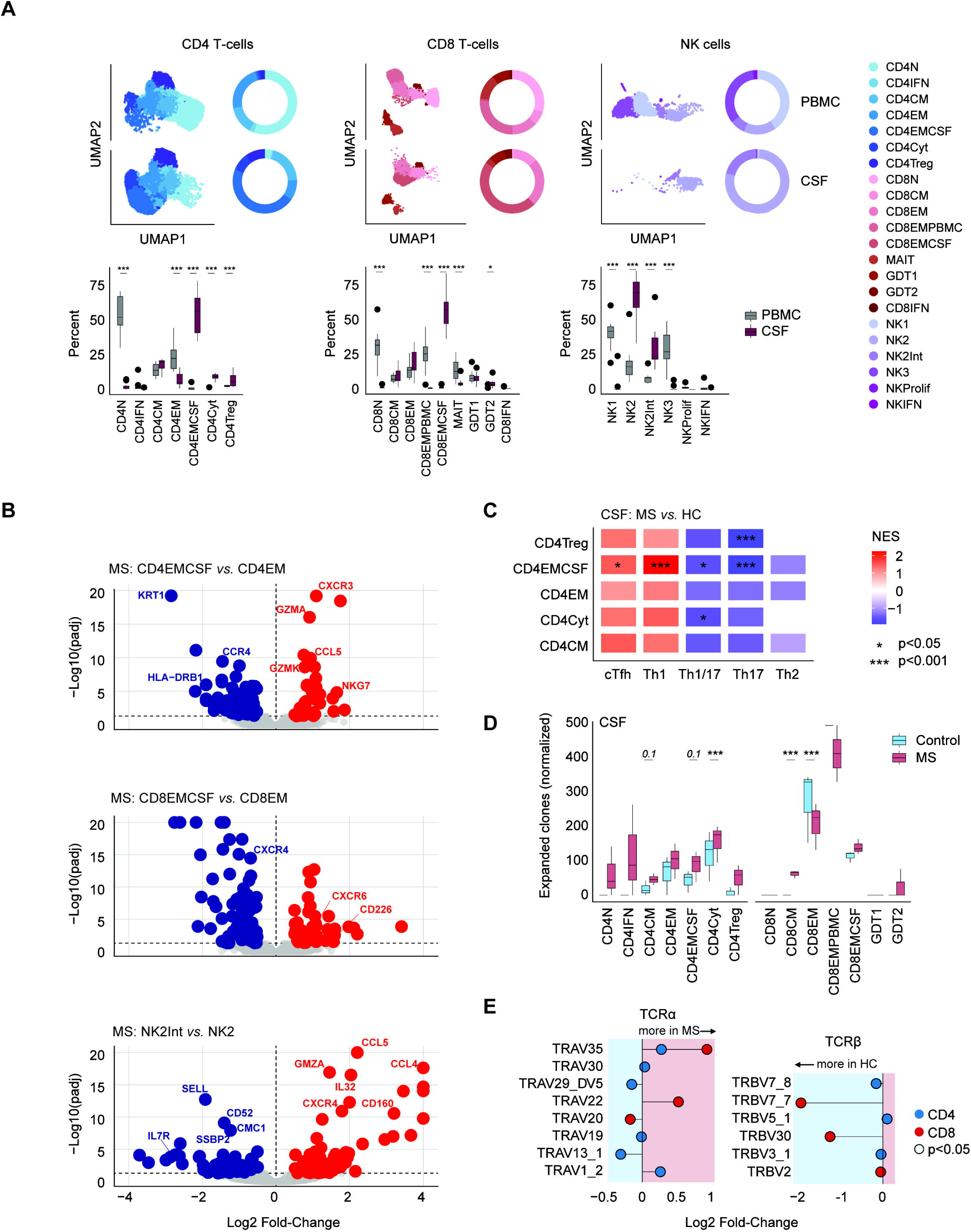
**A**. UMAP embeddings, donut plot of mean fraction of sub-clusters in each lineage, and boxplots of fraction of each sub-cluster in each lineage for CD4 T-cells (left), CD8 T-cells (middle) and NK cells (right), split by CSF and PBMC. **B**. Differential expression between shared and tissue-restricted effector memory subsets in people with MS: CD4EMCSF *vs.* CD4EM (top), CD8EMCSF *vs.* CD8EM (middle), NK2 *vs.* NK2int (bottom). Threshold for significance is indicated by dashed line. **C**. Visualization of enrichment scores after enrichment analysis (GSEA) of T helper (Th) signatures across disease in CD4 T-cells from CSF. **D**. Boxplots of normalized clonal number in CD4 (left) and CD8 (right) T-cell subsets in CSF, colored by disease status. Each point represents a donor. **E**. Relative log2 fold-change of variable genes on TCR A/B chains from V(D)J sequencing. Significant changes are outlined in black. *, p < 0.05; **, p < 0.01; ***, p < 0.001

To further characterize tissue-restricted counterparts, we performed differential expression analysis between CD4EMCSF *vs.* CD4EM, CD8EMCSF *vs.* CD8EM, and NK2 *vs.* NK2int in pwMS only (**Figure 2B**). In CD4 T-cells, tissue-restricted effector memory cells displayed significantly higher expression of cytotoxicity genes (*GNLY*, *GMZA/K/M*, *NKG7*, *PRF1*) and brain-homing *CXCR3* and *CCL5* genes. In contrast, the shared counterparts expressed HLA class II molecules, which typically associate with recent T-cell activation, and genes with immunoregulatory properties (*CREM*, *TIGIT, CTLA4*). Like CD4, the tissue-restricted NK2int population had increased expression of cytotoxic genes (*GRMA/M/H* and *NKG7*) and tissue-homing *CCL5*. In contrast, tissue-restricted CD8 effector memory T-cells expressed less of a defined phenotype, but shared with their CD4 and NK counterparts, increased expression of brain-homing molecules, such as *CXCR3* and *CXCR6*.

We next wanted to characterize CD4 T helper polarization in the CSF of pwMS, however, we found poor correspondence of sub-clusters to helper-subsets. We therefore performed Gene Set Enrichment Analysis (GSEA) on differential expression across disease in each sub-cluster, to understand the potential changes that are associated with MS. We observed a significant enrichment of Th1 and Tfh signatures and, interestingly, depletion of Th17 and Th1/17 signatures in CD4EMCSF of pwMS (**Figure 2C**).

To further characterize CD4 and CD8 T-cell responses in the CSF, we analyzed TCR sequences and defined expanded clones as similar TCRs based on amino-acid sequence in ≥ 2 cells. We observed an overall significant increase in the number of expanded clones across CD4 subsets in pwMS, with statistically significant increase in cytotoxic CD4 cells (CD4Cyt) and CD8CM, and a trend in CD4EMCSF, CD4CM and CD8EMCSF (p = 0.11, p = 0.11 and p = 0.11, respectively) cells (**Figure 2D**). Interestingly, there was a significant although modest decrease (β = –0.06) in the number of expanded CD8EM cells in pwMS. Since there was much individual variability in CDR3 regions, we sought to characterize variable gene usage. Indeed, we detected genes on both the alpha– and beta-chain that were dysregulated across disease in both CD4 and CD8 T-cells (**Figure 2E**). In pwMS, we found higher expression of *TRAV1-2*, *TRAV-30* and *TRBV5-1* in CD4, *TRAV22* in CD8, and *TRAV35* in both CD4 and CD8 T-cells.

Our data show a decrease in the relative abundance of tissue-restricted effector CD4 and CD8 T-cells in the CSF of pwMS. Similar to previous reports^18^, these cells are characterized by increased expression of cytotoxicity genes compared to their blood counterparts. Furthermore, we observed evidence of modulation of TCR signaling with enrichment of specific variable-genes, and increased number of expanded CD4 T-cell clones in the CSF of pwMS.

### IGHM-positive activated and atypical B-cells dominate CSF of pwMS

Similar to T-cells, naive B-cells (BN) dominate the lineage in blood, while activated (BAct) and atypical (BAtyp) B-cells represent the majority in the CSF (**Figure 3A, Supplementary Figure 5**). However, the CSF compartment is over-represented by B-cells derived from pwMS (**Figure 3B**). As previously reported^13–16,19,22^, only very few B-cells were observed in the CSF samples from HC (total 17 cells, mean 4.25), suggesting that BAct and BAtyp cells in the CSF are a hallmark of MS. We further validated that all B-cell subtypes are found in increased absolute numbers in pwMS as compared to matched healthy controls (**Figure 3C**).

**Figure 3.**
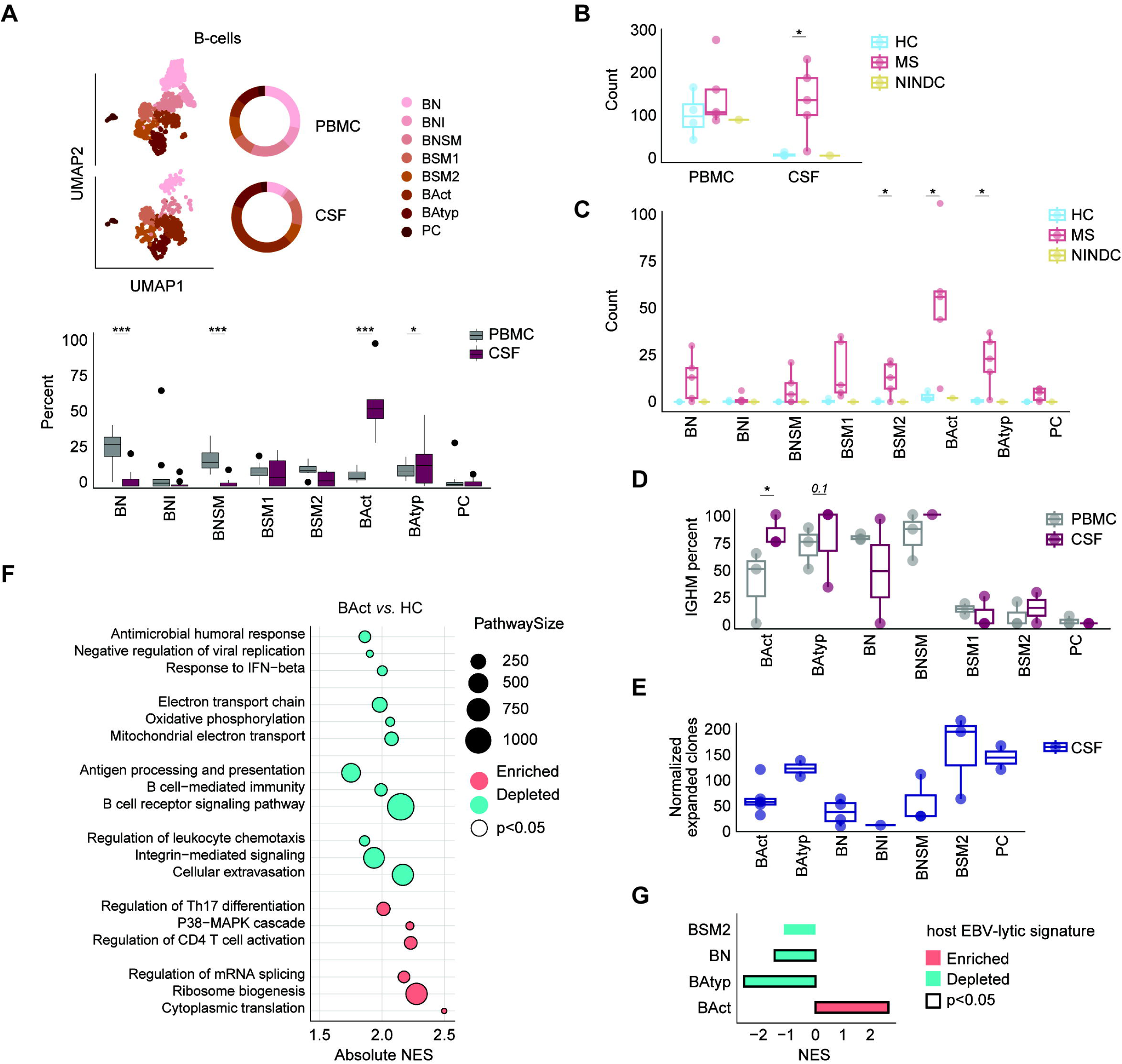
**A**. UMAP embeddings of B-cell sub-clusters (top left), donut plot of mean fraction of sub-clusters in B-cell lineage (top right), and boxplots of fraction of sub-clusters in B-cell lineage for each donor (bottom), split by CSF and PBMC. **B.** Boxplots of number of B-cells per sample. **C.** Boxplots of number of B-cells per sub-cluster. **D.** Proportion of B-cells with an IGHM-constant chain in each sub-cluster. **E.** Number of expanded clones per B-cell sub-cluster, normalized per 1000 cells. **F.** Selected biological processes from gene set enrichment analysis (GSEA) comparing activated B-cells with the rest. **G.** Gene set enrichment of the EBV-lytic gene signature across B-cell subsets in CSF of people with MS. *, p < 0.05; **, p < 0.01; ***, p < 0.001

Due to extremely low B-cell numbers in healthy individuals, we compared B-cell subpopulations between the periphery and CSF in pwMS only. We observed changes in constant chain usage, which reflects several key events during maturation^23^, with significantly more IGHM^+^ B-cells among BAct in the CSF compared to blood of pwMS (**Figure 3D**) and a trend of enrichment in BAtyp (p=0.1). Moreover, while BAct and BAtyp had comparable frequencies of IGHM^+^ cells as the naive B-cells in CSF of pwMS, they displayed a larger expanded pool of cells (**Figure 3E**). A GSEA on BAct *vs.* rest demonstrated significant depletion of pathways related to B-cell functions, energy and anti-viral responses, while CD4 T-cell activation, differentiation and protein synthesis were enriched in BAct-cells (**Figure 3F**). Moreover, BAct-cells in pwMS demonstrated the highest enrichment score of the lytic EBV transcriptomic signature^24^ across B-cell subsets, while the lytic EBV signature was depleted in BAtyp cells (**Figure 3G**).

Our findings validate previous reports^13–19^ that infiltration of B-cells into the CSF is a hallmark of MS and additionally demonstrate that they are predominantly non-class switched IGHM^+^ activated and atypical B-cells with expanded clonal BCRs, thus extending previous work^19^. We moreover report an enrichment for the EBV-lytic signature in activated B-cell subsets of pwMS.

### Increase of DCs and reduction of myeloid cells expressing markers of homeostatic microglia characterize CSF of pwMS

We approached analysis of sub-clustering of myeloid cells in a similar fashion, observing differences between the periphery and CSF (**Figure 4A, Supplementary Figure 6**). As expected CD14 and, to a lesser extent, CD16 monocytes made up the majority of the myeloid lineage in the periphery. The composition in the CSF was much more heterogenous. Previous work described a microglia-like myeloid cluster^19^, microglia^13,25^, CSF-resident macrophages with microglial-features^20^, and a disease-associated monocyte cluster^26^. We observed shared features with these clusters and annotated two microglia-like clusters (**Supplementary Figure 6**), one at resting state (MG-like), and one marked by higher expression of macrophage and activation markers (MG/MACa). MG-like and dendritic cells (DCs) were the most abundant subsets, followed by equal representation of CD14 monocytes, plasmacytoid dendritic cells (pDCs) and MG/MACa (**Figure 4A**). Within the myeloid compartment, we observed a significant enrichment of pDCs and a trend for enrichment of DCs and depletion of both MG-like and MG/MACa in the CSF of pwMS (**Figure 4B**). Due to limited cell numbers, we were restricted to comparing only MG-like cells in the CSF across disease. HLA class II genes, such as *HLA-DQA1*, *HLA-DPA*, *HLA-DQ1* and *HLA-DRB1* were increased in pwMS (**Figure 4C**), further supported by enrichment of antigen presentation and complement activation pathways in MG-like cells (**Figure 4D**).

**Figure 4.**
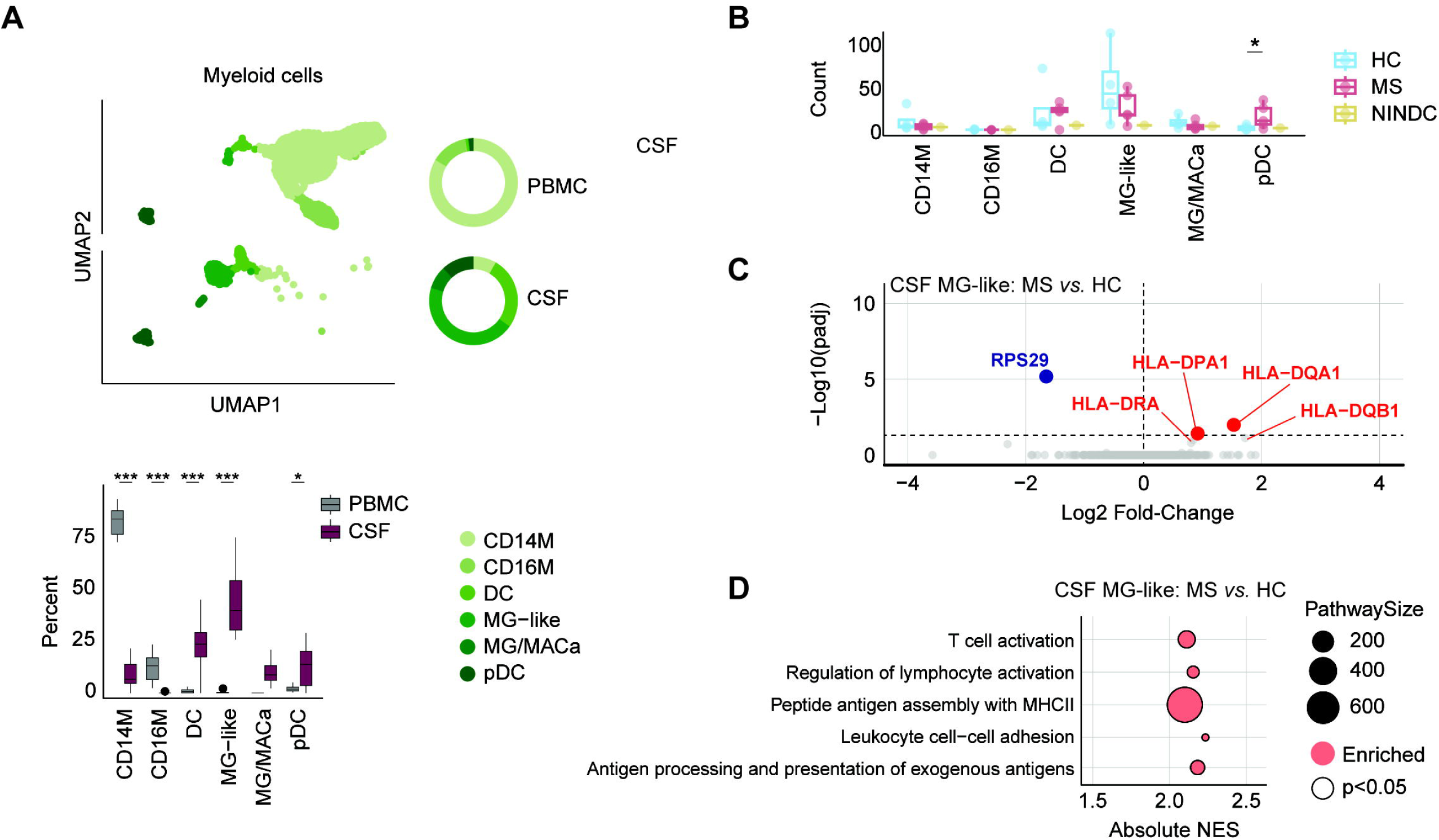
**A**. UMAP embeddings of myeloid sub-clusters (top left), donut plot of mean fraction of sub-clusters in myeloid lineage (top right), and boxplots of fraction of sub-clusters in myeloid lineage for each donor (bottom), split by CSF and PBMC. **B**. Boxplots of number of myeloid cell subsets per sample in the CSF. **C**. Differential expression between HC and MS, in the CSF MG-like cells. Threshold for significance is indicated by dashed line. **D**. Top biological processes from gene set enrichment analysis (GSEA) comparing MG-like cells across disease. *, p < 0.05; **, p < 0.01; ***, p < 0.001

We observed increased DCs and a trend of decreased microglia-like clusters in the CSF of pwMS, now compared to matched healthy subjects, replicating previous reports^13,19,20^. We also detected myeloid clusters expressing markers of microglia^19,20^, with MG-like cells from pwMS being characterized by features of increased antigen presenting functionality, suggesting that the CSF compartment may reflect the inflammatory milieu of the CNS tissue in pwMS.

## Discussion

Single-cell profiling of paired CSF and peripheral blood revealed coordinated and compartment-restricted alterations in cellular abundance and transcriptional signatures across multiple immune lineages in early MS. Activated and effector memory populations were enriched in the CSF of pwMS, whereas several of their counterparts were depleted from peripheral circulation, supporting the view that early MS is characterized by coordinated trafficking, activation and compartmentalization of immune cells to the CNS. This pattern was broadly replicated across several lineages, collectively suggesting that immune dysregulation in MS reflects dynamic interactions between the periphery and CNS, rather than purely local inflammatory events.

Multiple studies investigated inflammatory-milieu with single-cell resolution, predominantly comparing pwMS with other neurological diseases^13–20,27^ Our data validate a number of previous findings, including a significant enrichment of B-cells, memory T-cells, displaying Tfh features, Tregs and DCs in the CSF of early untreated MS compared to age– and sex-matched healthy controls. We further confirm previous findings of myeloid clusters expressing microglia genes^13,21^, which we find exclusively in the CSF though in low numbers. Utilizing freshly processed blood and CSF samples and matched healthy controls, we could extend these findings to further investigate paired changes with increased granularity.

We demonstrate a consistent pattern of discordant enrichment-depletion between periphery and CSF affecting multiple T and NK cell subsets. Tregs displayed particularly robust pattern and were, as previously reported^14,16,19^, significantly enriched in CSF, but concomitantly depleted from blood of pwMS compared to healthy controls. We observed a similar pattern for an NK subset that we designated as NK2, applying a more granular annotation from Rebuffet *et al*^18^. These NK2 cells expressed high levels of *CD56* and multiple migratory molecules, likely representing additional regulatory cell type suggested to be recruited to the CSF of pwMS^28^. The same discordant pattern was also observed across effector and central memory CD4 and CD8 T-cell subsets, strongly suggesting their recruitment from blood to CSF in early disease. Moreover, both CD4 and CD8 T-cells displayed a significant clonal expansion in CSF, but not blood, in pwMS compared to healthy controls. Similar to previous reports^27^, CD8 displayed more potent expansion compared to CD4 T-cells. Additionally, we detected altered expression levels of several TCR variable-genes on the clonal level. These changes in CD4 T-cells included overexpression of a variant of the *TRBV5* gene, previously detected in MS lesions^29^, as well as *TRAV35*, which was enriched in both CD4 and CD8 in pwMS. Interestingly, several variable genes were depleted in T-cells of pwMS, implying preferential variable gene usage. However, understanding which clonotypes are actually perturbed is difficult since the abundance of each clonotype may affect the relative proportion of the others.

We further characterized CSF-restricted lymphocyte subsets, including effector memory CD4 and CD8 T-cells (CD4EMCSF and CD8EMCSF) and NK cells (NK2Int), that were all significantly depleted from the CSF of pwMS. Interestingly, all three subsets displayed more pronounced cytotoxic profile compared to their effector counterparts that were present both in blood and CSF. Moreover, the CD4 CSF-restricted effectors were enriched in the Th1 signature genes in pwMS compared to healthy controls, and may explain the reported decrease of Th1 cells in CSF of pwMS in the recent integrative analysis by Cantoni *et al*^21^. Interestingly, these cells were characterized by a decreased profile of Th17, and to a lesser extent Th1/17, signature genes. It is plausible that these tissue-primed proinflammatory and cytotoxic effector cells get recruited to the CNS tissue where they exert pathogenic effects, which is supported by their higher expression of brain-homing molecules such as *CXCR3*, *CXCR6* and *CCL5*.

A hallmark of MS is the accumulation of B-cells in the CSF, which was also prominent in our cohort, where only a handful of B-cells were captured in the CSF of healthy individuals. Cantoni *et al.*^21^ reported dramatic expansion of atypical memory B-cells, and we could further show that the CSF compartment of pwMS is dominated by expanded IGHM-positive activated and atypical B-cells. We found that activated B-cells were enriched in pathways related to CD4 T-cell activation and differentiation, suggesting that they may be involved in mediating the B-T interactions in MS. Interestingly, they were also depleted in pathways related to B-cell activation, function and anti-microbial response and displayed a significant enrichment of a “host lytic EBV– related transcriptional signature”^31,32^ B-cells infected with EBV have been detected in the CSF of healthy individuals and pwMS^31^ yet it is still unclear which B-cells are targeted. Our data suggest that activated B-cells potentially engage in the initial B-T interactions following either EBV infection or reactivation. However, the “host lytic EBV-related transcriptional signature” is dominated by protein replication machinery, suggesting that we are likely assaying transcriptional burden of B-cells, which may or may not reflect the EBV infection state. Recent *in-vitro* characterization of auto-proliferating B-cells from pwMS demonstrated their shared features with atypical age-associated B-cells (ABC), the potential for extra-follicular B-T cell interactions and established IGHM-positive clonotypes as the most abundant atypical B-cell type^9^. Accumulation of autoimmune atypical B-cells outside the B-cell follicle has been reported^33^ and it was proposed in lupus that human ABCs may be generated predominantly through an extrafollicular response^34^. Thus, our findings further imply the role of IGHM^+^ atypical B-cells in extra-follicular B-T cell interactions in pwMS *in-vivo*.

We recognize that there are several limitations to our study. Firstly, despite analyzing fairly homogeneous and matched groups, we are limited by the small number of participants, especially when it comes to analyzing TCR and BCR sequences, posing a risk for an inherent bias. Secondly, we are technically limited by the number of analyzed cells, specifically from CSF, and estimates obtained from cell subsets with small numbers may be inherently noisy. Finally, we recognize that sequencing technologies capture a single-snap shot of cell states, which should be recognized when interpreting results. Integrated analyses of multiple datasets across disease trajectory and conditions will be crucial for characterizing the CSF immunoenvironment in MS.

Our data highlight MS as a highly compartmentalized immune disorder characterized by dynamic interactions between periphery and CNS. We provide a detailed characterization of the immune landscape of early MS in blood and CSF, and a reference for validation of future work in matched healthy controls.

## Statements and Declarations

### Ethical Considerations

Ethical approval (2009/2107-31, last amendment 2025-04961-02), was obtained from the Swedish Ethical Review Authority in Stockholm, Sweden (Regionala etiksprovnämden I Stockholm).

### Consent to participate

All participants in this study signed consent forms.

### Consent for publication

Not applicable.

### GenAI usage disclosure

Generative AI was limited in use to resolving errors during analysis and help with generating certain helper functions, which were functionally validated.

