## Supplementary Methods for "Single-cell analysis suggests coordinated immune-cell redistribution between blood and cerebrospinal fluid in early multiple sclerosis"

##### Sample collection

Study participants were recruited at the Neurology clinic at Karolinska University Hospital in Stockholm, Sweden. They were newly diagnosed with MS, with no previous treatment history. Healthy controls were age- and sex-matched. All study participants provided written consent.

Mononuclear cells were isolated from peripheral blood using a standard Ficoll method. CSF cells were centrifuged at 300 RCF for 10 minutes, re-suspending in 2mL of PBS, and re-centrifuged for another 10 minutes at 300 RCF.

##### Library Preparation

Freshly isolated cells from PBMC and CSF were loaded onto Chromium Single Cell Controller using the Chromium Single Cell 5' library and gel bead kit (Cat. No. PN-1000006 and PN-10000014). Expression and VD(J) library amplification was performed on the Chromium Single Cell 5' Library Construction Kit, (Cat. No. PN-10000020), Chromium Single Cell V(D)J Enrichment Kit for Human B Cell (Cat. No. PN-10000016) and Chromium Single Cell V(D)J Enrichment Kit for Human T Cell (Cat. No. PN-1000005), and Chromium i7 Multiplex Kit (Cat. No. PN-120262) as per manufacturers instructions. Sequencing was carried out on the Illumina Next Seq and Miseq platforms with read length per 10X protocol. Cell Ranger pipeline v 3.1.0 (10X Genomics) was used to process reads and align to GRCh38 genome reference.

##### Pre-processing and quality control

To address ambient contamination of mRNA, we processed the unfiltered output from *CellRanger3* using *CellBender* on each sample. *CellBender* uses a generative model to approximate latent variables and generate a flexible prior for inference of a denoised-count matrix with a Bayesian framework. Default parameters were passed, and *CellBender* was allowed to estimate the total number of cells and expected droplets automatically. Samples were loaded into R (v4.4) for preliminary QC analysis (**Supplementary Figure 1**). Low-quality cells were excluded from the dataset per-sample basis by excluding cells that exhibited nUMIs or nGenes outside of 3 median absolute deviations. Furthermore, cells that showed a fraction of mitochondrial reads greater than 15%, presumably due to damage and/or death, were also removed from the dataset.

Droplet-based assays are susceptible to multiplets, droplets that contain more than one cell and can introduce undesirable results in downstream analysis. The package *DoubletFinder* was used to remove droplets. Briefly, *DoubletFinder* uses clustering annotation to generate synthetic doublets of pairs of cells from different clusters. Real cells proximal to the synthetic cells are classified as doublets. On a per-sample basis, mitochondrial and ribosomal genes were removed from the count matrix (genes starting with MT-|RPS|RPL). The count matrix was normalized and log-transformed with *Seurat*'s NormalizeData function (default parameters). HVGs were selected using *Seurat*'s FindVariableFeatures function (selection method = 'vst'). Counts were scaled, and dimensional reduction (PCA, dims 1:20, UMAP) was run as implemented in *Seurat*. Estimation of the pK value for *DoubletFinder* was performed automatically at each point. The maximal pK value from paramSweep was used in the *DoubletFinder* function with parameters 20PCs, and an estimated droplet frequency of ~2%. Cells identified as doublets were removed. Matrices from each sample were aggregated and exported for downstream analysis. Two donors were excluded from the downstream analysis. In one donor sample, one of the tissue pairs did not sequence properly. The second donor exhibited strong batch effects, which made integration with the data difficult.

*List of packages used: Seurat, DoubletFinder, anndataR, ggplot2, CellBender*

**Preprocessing of TCRs**

TCRs were initially loaded from the 10X all_contigs.json. Unproductive and multi/orphan-receptors were removed. A pairwise distance matrix was calculated between each alpha and beta chain, scoring for amino-acid similarity based on BLOSUM62 as implemented in scripy. The default cut-off was used to determine distances for clonotype clusters. We defined clonal groups as expanded, if there were ≥ 2 cells within a clonotype cluster.

*List of packages used: scirpy*

**Preprocessing of BCRs**

BCRs were initially loaded from the 10X fasta file output. The package *dandelion* was used to re-annotate BCR-genes with default settings and assign isotopes, using the *IgBlast* backend. The *dandelion* object was directly exported to *scirpy*, where unproductive BCR chains and orphaned/multichain/ambiguous V(D)J chains were removed. To calculate clonality of BCR clones, we calculated a pair-wise distance matrix between each of the heavy and light chains, using a normalized hamming distance on the nucleotide sequence as implemented in *scirpy*. We defined clonal groups as expanded, if there were ≥ 2 cells within a clonotype cluster.

*List of packages used: dandelion, IgBlast, scirpy*

##### Initial clustering of all lineages

The *Anndata* object was loaded into *scanpy*. Both cells from PBMC and CSF were clustered together, as well as cells from healthy controls and pwMS. Prior to the selection of highest-variable genes (HVGs), mitochondrial and ribosomal genes were removed from the matrix. Furthermore, the gene *MALAT1* was also excluded, as previous studies have reported that *MALAT1* may report intron load. The denoised count matrix was normalized to library size, and log transformed with a pseudocount of +1, implemented in *scanpy* with default parameters. The selection of HVGs was performed using *scanpy*'s implementation of *Seurat* version3; the top 2000 genes were selected, passing the sequencing runs as the argument to the batch-key parameter. Counts were unit-scaled to enforce equal weights of transcripts in dimensional reduction tasks with *scanpy*'s pp.scale function, clipping values at 10. Principal component analysis (PCA) was performed to reduce complexity prior to constructing a nearest-neighborhood graph. PCA-embeddings were corrected with *Harmony* across individuals and was run until convergence. We constructed a nearest-neighborhood graph from corrected PCs and UMAP embeddings with *scanpy*. Cells were clustered into discrete partitions using the leiden algorithm implemented in *scanpy* (**Supplemental Figure 1**).

*List of packages used: Scanpy, Anndata, harmonypy, muon*

##### Sub-clustering of each lineage

We sub-clustered each lineage in a similar fashion as the parent analysis. Denoised counts for cells from each coarse lineage were loaded in, and nuisance genes were removed from the count matrix. The counts were normalized, log-transformed, and scaled, as described in the previous clustering section. Embeddings for the dataset were generated with PCA and integrated with *Harmony*. The selection of PCs for each sub-cluster was made somewhat arbitrarily; the number of PCs was chosen at the inflection of the elbow plot. Clustering resolution differed based on lineage type, where a resolution was chosen based on expected cell-types.

Contamination of T-cells was an issue in myeloid, B-cell and NK-cell sub-clustering. In some clusters we observed T-cell specific transcripts. In such cases, we validated with presence of TCRs and discrete clusters that seemed to be driven by T-cell contamination were removed, and the sub-clustering was performed again.

*List of packages used: Scanpy, Anndata, harmonypy, muon*

**Annotation of cluster identities**

Seven discrete clusters of CD4 T-cells (**Supplementary Figure 2**) were identified and annotated based on canonical markers into: naive CD4 (**CD4N**; *SELL*, *TCF7*, *CCR7*, *LEF1*), naive-like IFN responding CD4 (**CD4IFN**; *ISG15*), central memory (**CD4CM**; *TCF7*, *CCR7*, *PASK*), effector memory (**CD4EM**; *IL7R*, *KLRB1*/*CD161*), tissue-specific EM (**CD4EMCSF**; *IL7R*, *KLRB1*/*CD161*, *GNLY*), cytotoxic CD4 (**CD4Cyt**; *GNLY*, *PRF1*, *CXCR3*, *CCL5*, *CST7*), and regulatory T cells (**CD4Treg**; *FOXP3*, *TIGIT*).

Nine discrete clusters of CD8 T-cells (**Supplemental Figure 3**) were identified and annotated based on canonical markers into: naive CD8 (**CD8N**; *SELL*, *TCF7*, *CCR7*) and a naive-like cluster expressing IFN-related genes (**CD8IFN**; *ISG15*, *MX2*), central memory (**CD8CM**; *SELL*, *TCF7*, *CCR7*, *TRAC*, *ITGB1*, *CD27*, *GPR183*) and three effector memory subsets based on loss of stem markers (*SELL*, *TCF7*, *CCR7*, *LEF1*), one primarily found in blood (**CD8EMPBMC**), one primarily found in CSF (**CD8EMCSF**), and a shared effector subset between the two tissues (**CD8EM**).

Six discrete clusters of NK-cells (**Supplemental Figure 4**) were identified and annotated based on sub-populations previously identified in blood (Rebuffet L, *et al. Nat Immunol.* 2024 https://doi.org/10.1038/s41590-024-01883-0) into: NK-cells with cytotoxic functions (**NK1**; *GZMB*, *PRF1*, *CLIC3*), expression of migratory receptors (**NK2**; *CD56/NCAM1*, *CD44*, *SELL*, *GPR183*, *GZMK*), resembling adaptive NK cells (**NK3**; *CCL5*, *CD3E*, *IL32*, *GZMH*), an intermediate NK2-cluster (**NK2Int**; *ZFP36*, *CXCR4*, *IER2*), IFN-activated (**NKIFN**; *ISG15*, *CD69*) or proliferating NK cells (**NKProfil**; *TOP2A*, *PCNA*).

Eight discrete clusters of B-cells (**Supplemental Figure 5**) were identified and annotated based on canonical markers into: naive (**BN**; *SELL*, *TCL1A*, *IGHM*, *IGHD*), non-switched B memory (**BNSM**; *IGHM*lo, *IGHD*lo, *CD27*), two clusters of switched B memory (**BSM1** and **BSM2**; *IGHM*neg, *IGHD*neg, *CD27*), IFN-stimulated B memory (**BNI**; *ISG15*, *MX1*), atypical B cells

(**BAtyp**; *FCLR5*, *TBX21*, *CXCR3*), activated B cells (**BAct**; *CD83*, *JUN*) and plasma cells (**PC**; *CD38*, *JCHAIN*, *MZB1*).

Eight discrete clusters of myeloid cells were identified and re-annotated as six different cell types (**Supplemental Figure 6**) based on canonical markers into: CD14 monocytes (**CD14M**; *CD14*, *VCAN*, *FCN1*, *LYZ*, *S100A8/9*, clusters 0-2), CD16 monocytes (**CD16M**; *CD16a/FCGR3A*, *CD11a/ITGAL*, *LST1*, cluster 3), a heterogeneous group of conventional dendritic cells expressing both cDC1 and cDC2 markers (**DC**; *CDC1C*, *FCER1A*, *CLEC10A*, *XCR1*, cluster 5), a myeloid cluster expressing markers of homeostatic microglia (**MG-like**; *TMEM119*, *P2RY12*, *C1QA*, cluster 4), plasmacytoid dendritic cells (**pDC**; *LILRA4*, *TCF4*, *GZMB*, cluster 6) and a myeloid cluster expressing macrophage and activation markers including markers of disease-associated microglia (**MG/MACa**; *CCL3*, *CD63*, *CD9*, *APOE*, *TREM2*, *IL1B*, cluster 7).

**Differential expression**

Cells in each lineage by donor were sum-pseudobulked using *decoupler* on ambient-mRNA denoised counts. Low-count genes were removed from the matrix. Samples were removed if they had less than 20 cells, or less than 1000 total counts. For each comparison, if the number of samples was < 3 that did not meet the QC cutoff, it was skipped. *Pydeseq2* was used to fit a negative-binomial generalized-linear model for each gene. For cluster-level analysis, the model fit was y ~ cell-type + individual, whereas for cell-type-specific disease analysis, the model fit was y ~ disease + batch. P-values for each test are automatically adjusted for multiple-testing correction with Benjamini-Hochberg, and genes whose q-value < 0.05, were considered significant for downstream visualization and reporting.

*List of packages used: decoupler, pydeseq2, Anndata, muon*

**Differential abundance**

For each cell type, we modelled cell counts as a proportion of the total cells using a beta-binomial model from the *VGAM* package. Comparisons where there were less than three non-zero fractions in a group were excluded. Log2 fold-change was derived from ratio of group-level mean predicted proportions. Multiple-testing was corrected for with Benajamini-Hochberg.

For differential abundance of TCRs we implemented a similar approach, where the primary annotation for each v-gene from the alpha- and beta-chain was used. We aggregated per group the sum of number of cells which contained each gene. Total number of cells with productive

TCRs were used as the denominator instead of total cells. Like cell abundance, comparisons where there were less than three non-zero factions in a group were excluded. Only samples where VDJ sequencing was performed were considered.

*List of packages used in the differential abundance step: VGAM, r2py*

**Gene Set Enrichment**

A ranked gene-list was created from the T-statistics after differential expression. GSEA was performed as implemented in *decoupler*. Gene-set annotation pathways were downloaded from msigdb.

*List of packages used: Decoupler, msig*

**Discordant analysis**

Log2fold changes, beta-values and standard errors from abundance analysis were used. Since many of our estimates did not meet significance after multiple testing correction, we assumed that our beta-values may be noisy. We particularly care about the reliability of the sign (direction) of the beta value. We estimate the probability distribution that our beta-value reflects the true sign change. To understand whether we observe more discordant relationships than one would expect by arbitrary pairing, we perform a permutation test, assigning a random sign to one of the feature pairs, and a one-sided empirical p-value was calculated.

*List of packages used: scipy*

**Visualization**

All visualizations were performed in R with *ggplot2*.

#### Supplementary Figure Legends

**Supplementary Figure 1. A.** Boxplot of number of cells acquired per donor in PBMC (left) and CSF (right), colored by disease status. Each dot represents a donor. **B.** Violin plots of number of genes > 1 count (top row), total counts (second row), percent mitochondrial reads (third row), and percent ribosomal reads (fourth row) across each sample, colored by disease status. **C.** UMAP visualizations pre- and post-correction of PCA colored by individuals (top) and runs (bottom). **D.** UMAP visualization colored by initial clusters. **E.** Dot plot of canonical marker genes used to assign initial clusters into coarse lineages. **F.** Stacked bar of fraction of initial clusters from each donor. **G.** UMAP embeddings of initial clustering annotated by lineages.

**Supplementary Figure 2. A.** UMAP embeddings of CD4 T-cell sub-clustering, colored by annotated clusters. **B.** Dot plot of canonical marker genes for each of the annotated CD4 T-cell sub-clusters. **C.** Boxplots of total number of CD4 T-cells in PBMC (left) and CSF (right), colored by disease status. Each dot represents an individual donor**. D.** Stacked bar-plot of fraction of annotated CD4 T-cell clusters in each sample for PBMC (left) and CSF(right). **E.** Stacked bar-plot of fraction of CD4 T-cells which VDJ-sequences were obtained for in each sample.

**Supplementary Figure 3. A.** UMAP embeddings of CD8 T-cell sub-clustering, colored by annotated clusters. **B.** Dot plot of canonical marker genes for each of the annotated CD8 T-cell sub-clusters. **C.** Boxplots of total number of CD8 T-cells in PBMC (left) and CSF (right), colored by disease status. **D.** Stacked bar-plot of fraction of annotated CD8 clusters in each sample for PBMC (left) and CSF (right). **E.** Stacked bar-plot of fraction of CD8 T-cells which VDJ-sequences were obtained for in each sample.

**Supplementary Figure 4. A.** UMAP embeddings of NK cell sub-clustering, colored by annotated clusters. **B.** Dot plot of canonical marker genes for each of the annotated NK sub-clusters. **C.** Boxplots of total number of NK cells in PBMC (left) and CSF (right), colored by disease status. **D.** Stacked bar-plot of fraction of annotated NK clusters in each sample for PBMC (left) and CSF(right).

**Supplementary Figure 5. A.** UMAP embeddings of B-cell sub-clustering, colored by annotated clusters. **B.** Dot plot of canonical marker genes for each of the annotated B-cell sub-clusters. **C.** Stacked bar-plot of fraction of annotated B-cell clusters in each sample for PBMC (left) and CSF(right). **D.** Relative log2 fold-change of sub-cluster proportions in each tissue compartment across disease.

**Supplementary Figure 6. A.** UMAP embeddings of myeloid cell sub-clustering, colored by annotated clusters. **B.** Dot plot of canonical marker genes for each of the annotated myeloid sub-clusters. **C.** Boxplots of total number of myeloid cells in PBMC (left) and CSF (right), colored by disease status. **D.** Stacked bar-plot of fraction of annotated myeloid clusters in each sample for

PBMC (left) and CSF(right). **E.** Relative log2 fold-change of sub-cluster proportions in each tissue compartment across disease. Significant values are denoted with a black outline.

Supplementary Figure 1

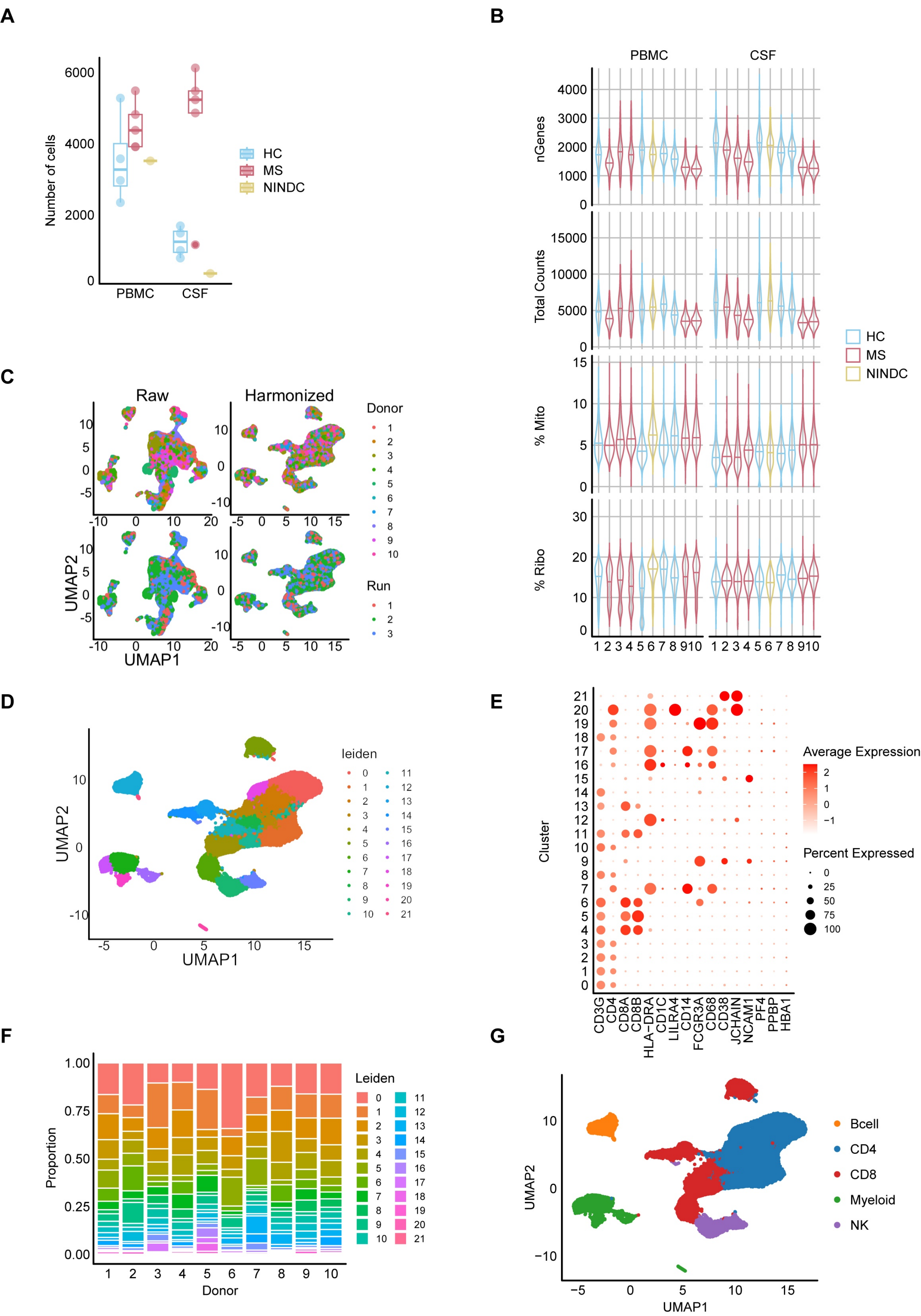

Supplementary Figure 2

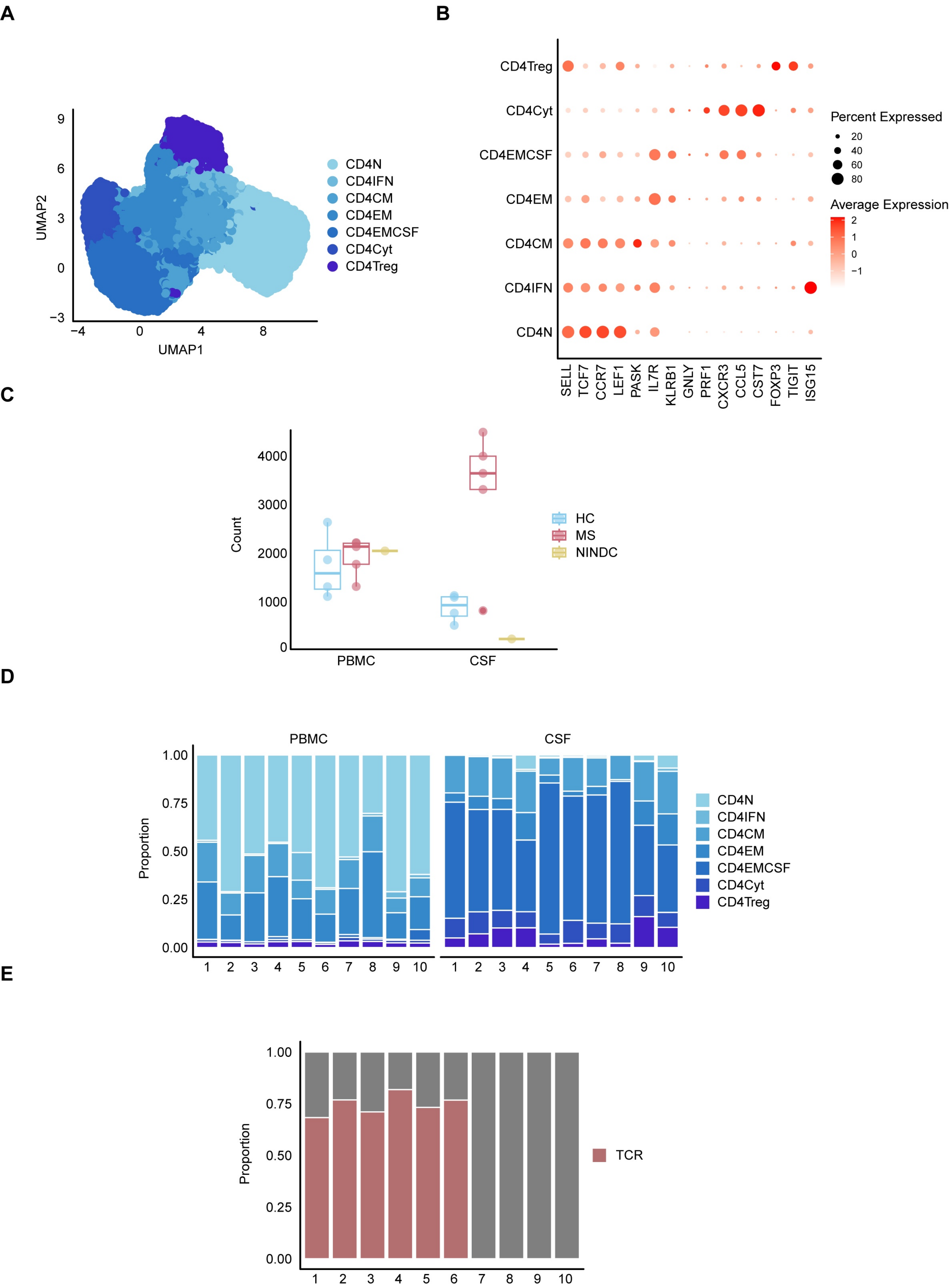

### Supplementary Figure 3

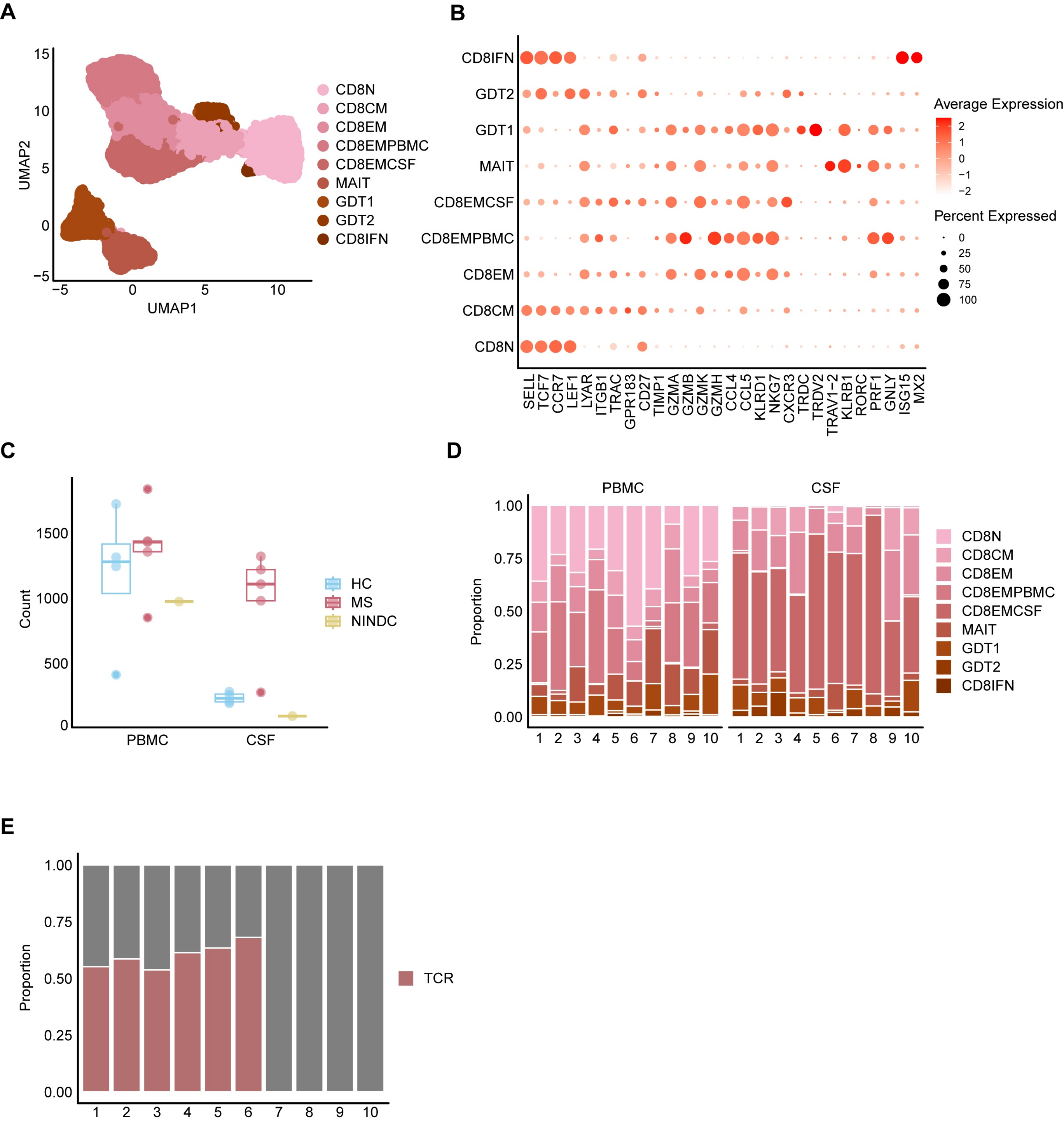

### Supplementary Figure 4

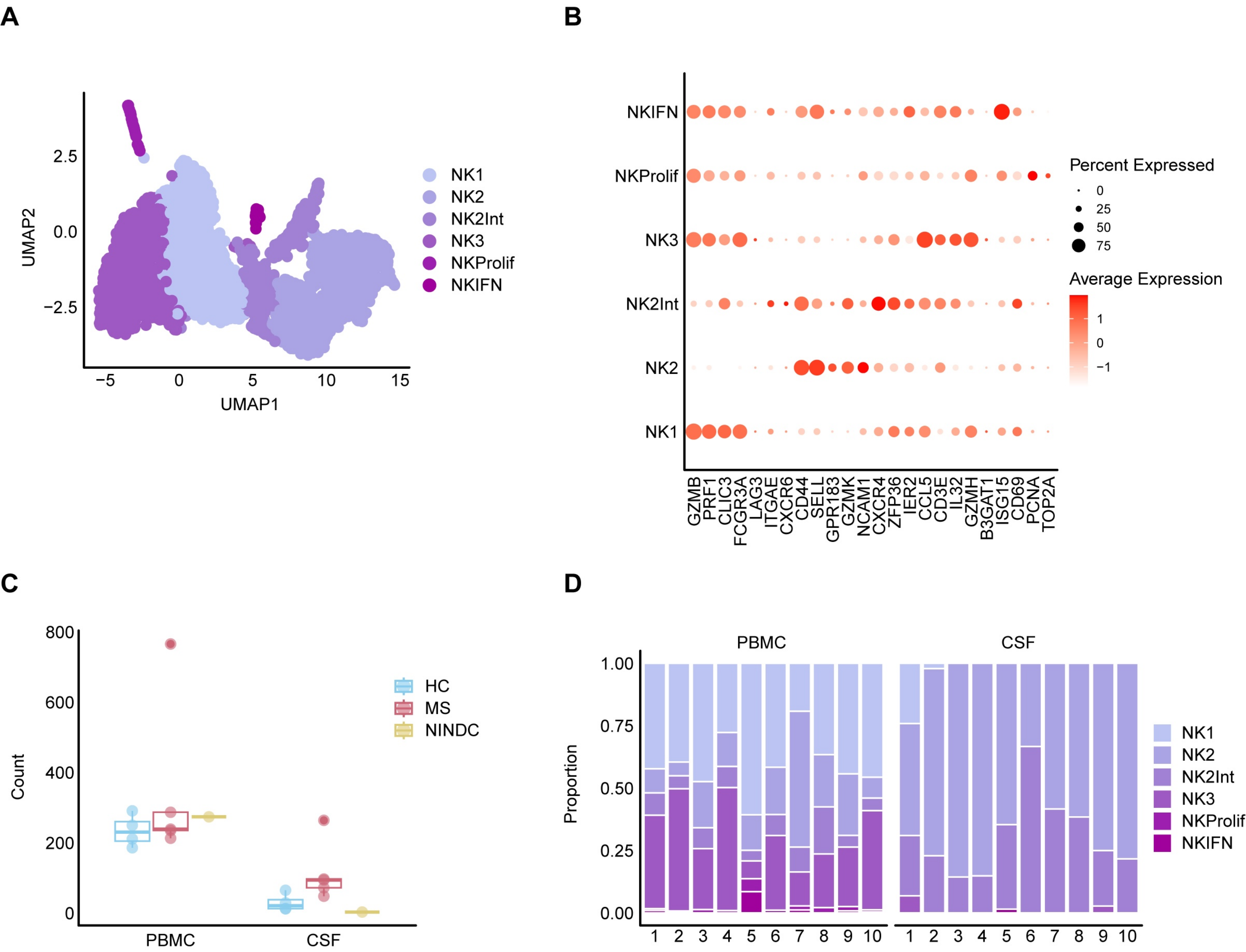

#### Supplementary Figure 5

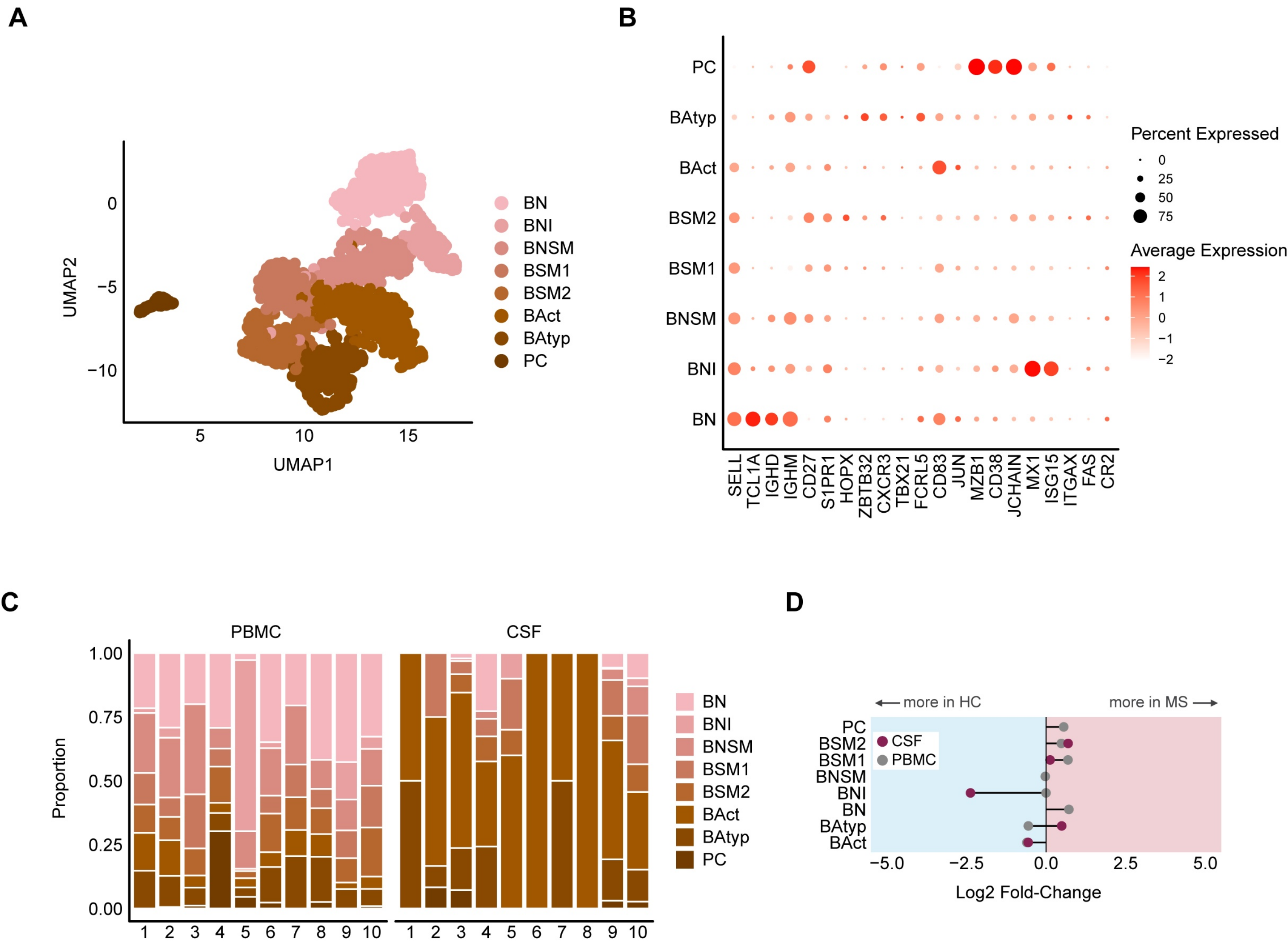

### Supplementary Figure 6

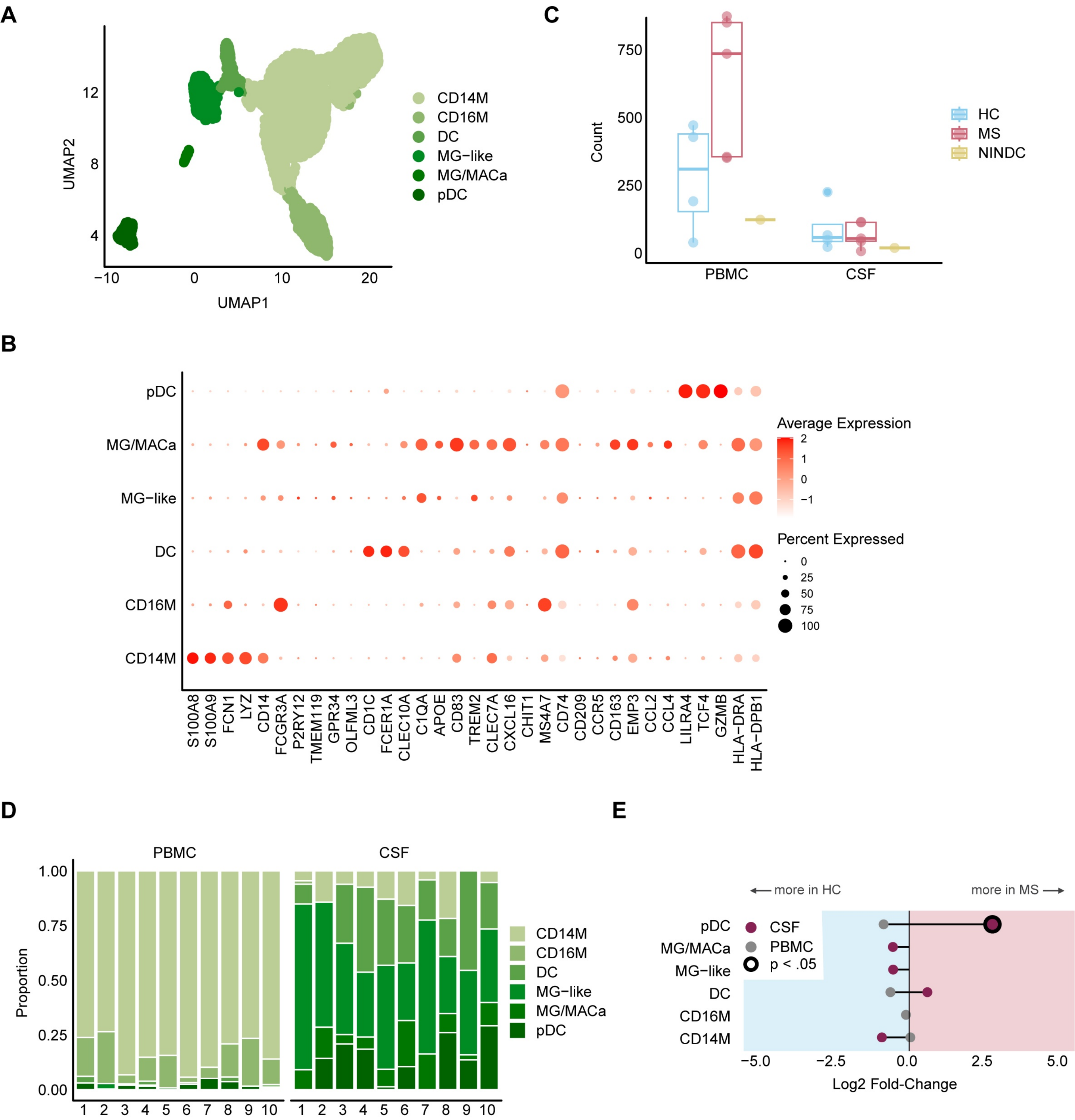
